# Centering based on active diffusion in mouse oocytes is non-specific

**DOI:** 10.1101/531657

**Authors:** Alexandra Colin, Nitzan Razin, Maria Almonacid, Wylie Ahmed, Timo Betz, Marie-Emilie Terret, Nir S Gov, Raphaël Voituriez, Zoher Gueroui, Marie-Hélène Verlhac

## Abstract

The mechanism for nucleus centering in mouse oocytes results from a gradient of actin-positive vesicles. By microinjecting oil droplets and fluorescent beads, we analyze the consequences of the gradient of activity on transport of exogenous tracer particles of different sizes. We also use optical tweezers to probe rheological properties of the cytoplasm. We find that the gradient activity induces a general centering force, akin to an effective pressure gradient, leading to centering of oil droplets with velocities comparable to nuclear ones. High temporal resolution measurements reveal that passive particles, larger than 1µm, experience the activity gradient by a biased diffusion towards the cell center. Unexpectedly, this general and size dependent mechanism is maintained in Meiosis I but contrasted by a further process that specifically off-centers the spindle. These antagonizing processes depend on myosin activity, thus we reconcile how the same molecular actors can have two opposite functions (centering versus off-centering).

## Introduction

The position of the nucleus in a cell can instruct morphogenesis, conveying spatial and temporal information. In addition, abnormal nuclear positioning can lead to disease^1^. In mammals, the oocyte nucleus is centered via actin-based mechanisms^2^. Importantly an off-centered nucleus correlates with poor outcomes for mouse and human oocyte development^3,4^. This is surprising since oocytes subsequently undergo two extremely asymmetric divisions in terms of size of the daughter cells, requiring an off-centering of their chromosomes^5–7^.

Using a multidisciplinary approach, we recently discovered how the nucleus is actively centered in mouse oocytes^2^. We observed that oocytes derived from Formin 2 (Fmn2) knockout mice, which lack actin filaments in their cytoplasm, present off-centered nuclei^6,8^. Formin 2 is a straight microfilament nucleator and also an essential maternal gene^9^. The re-introduction of Formin 2 into *Fmn2-/-* oocytes, that harbor initially off-centered nuclei, induces a directional motion of the nucleus toward the center within about 5 hours in 100 % of the cases^2^. We found evidence suggesting the existence of a centering force, akin to an effective pressure gradient, exerted by the actin mesh, that acts on the nucleus to move it from the periphery to the center^2^. In this model system, actin filaments are nucleated from Rab11-a positive vesicles since actin nucleators, such as Formin 2 and Spire 1&2 are anchored there^10^. We have shown that these actin-positive vesicles are biased in their speed, moving faster at locations closer to the cortex. Accordingly, the spontaneous fluctuations, or activity of actin-positive vesicles decreases from the cortex to the oocyte center as quantified by their squared velocity *v*^2^. On the basis of a simple model describing the pool of actin-positive vesicles as an ideal suspension of self-propelled particles, we suggested that this gradient of activity from actin vesicles, which move by active diffusion^2^, generates an effective pressure gradient^11–13^ and thus a propulsion force. It is thereby the trigger of the nuclear motion towards the oocyte center^14^. In addition, recent evidence has shown that active diffusion is also a major player of organelle motion in the cytoplasm of *Drosophila* oocytes^15^.

As this active-pressure centering mechanism should not be specific to the nucleus, we tested it by micro-injecting oil droplets as well as fluorescently-labeled latex beads of various sizes and monitoring the localization of these exogenous objects. This also allowed us to probe the spatiotemporal rheological properties of the actin cytoskeleton and analyze the transport properties of exogenous passive particles of different sizes and chemical nature. We report here, that nuclear sized, but fully passive oil droplets are centered with velocities comparable to the nuclear centering and that they phenocopy the nuclear envelope in terms of actin filament recruitment. This indicates that the gradient of activity does not require any specific signaling to move the oocyte nucleus and is able to center other objects in addition to the oocyte nucleus. At temporal resolution of seconds, we show that objects larger than 1 µm in diameter experience a significant biased movement toward the center of the oocyte, consistent with an active pressure gradient. Interestingly, the centering velocity over much longer time scales of hours seems to be independent of the particle size for larger objects such as oil droplets of diameters comprised between 5 to 35 µm.

In addition, a puzzling question in the field is how the same molecules, namely Formin 2, Spire 1 & 2 and Myosin Vb are able to promote two opposite motions: centering of chromosomes enclosed inside a nucleus in Prophase I^2^ and off-centering of chromosomes enclosed inside a meiotic spindle later in meiosis I^7,10,16–19^. We offer here an explanation to this long-standing question. Indeed, oil droplets injected in Meiosis I also undergo centering, in a cytoplasm that is mechanically comparable to Prophase I oocytes as measured by optical tweezers. Thus, a similar passive mechanism, able to non-specifically center organelles, does co-exist in Meiosis I together with a specific process that depends on Myosin II^16–18,20^ and which promotes spindle off centering. We reconcile here how the same molecular actors can be involved in two opposite functions (centering versus off-centering of objects).

## Results

### The centering mechanism is not specific to the biological nature of the nucleus

We used inert oil droplets as passive particles to test whether the centering mechanism is specific or not to the nucleus. For this purpose, we injected oil droplets in Prophase I mouse oocytes and observed them on short time scales (min). These oil droplets present 1.9 times the density of water and maintain a round shape indicating that the forces generated in the cytoplasm are much weaker than capillary forces.

We recorded movies with one image every 500 ms (Supplementary Video 1), and tracked the oil droplet centroid (Methods) to analyze the nature of the observed motion. First, to check for a potential bias in droplets’ motion, we measured the radial (∆r.cos(θ)) and the orthoradial (∆r.sin(θ)) components of the displacement vector of the droplet centroid over a time step Δt (Fig. 1a). For Δt = 500 ms we observed a statistically significant bias of the displacement vector towards the center of the cell in the presence of F-actin (Fig. 1b and 1c and Supplementary Fig. S1a). In contrast, in oocytes treated with Cytochalasin D (Supplementary Video 2), no statistically significant bias was observed, consistent with isotropic fluctuations inside the cell in the absence of F-actin (Fig. 1b, 1c, Supplementary Fig. 1a). This bias on the radial component was confirmed by analyzing the cumulative probability of negative values (Supplementary Fig. S1b). At long time scales (15-25 s), the cumulative radial bias toward the center increased in oocytes in Prophase I, whereas it stayed below the noise on the orthoradial component (Supplementary Fig. S1b). In oocytes treated with Cytochalasin D, cumulative bias on radial and orthoradial components stayed constant at all time scales (Supplementary Fig. S1b).

**Figure 1.**
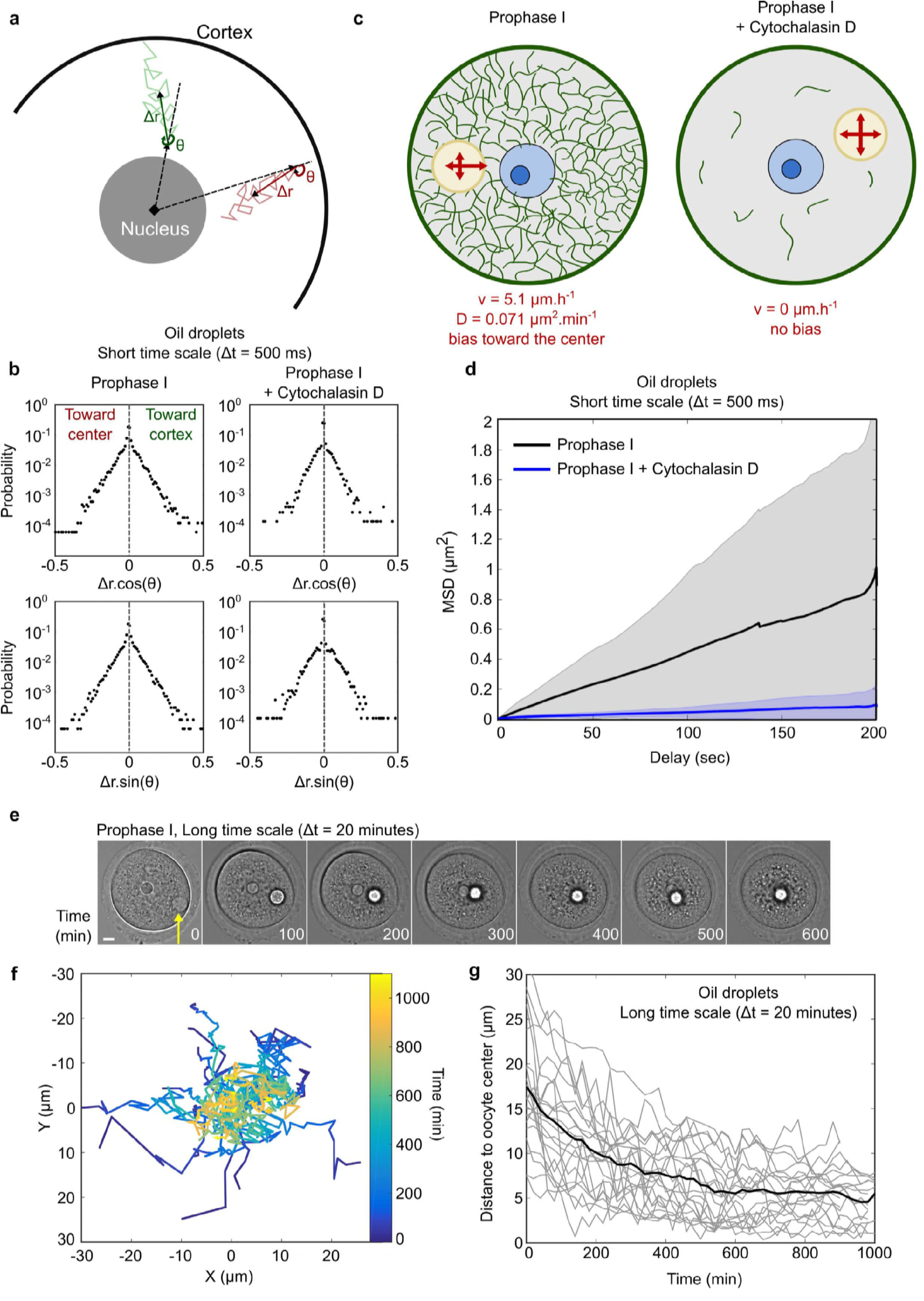
Peripherally injected oil droplets are centered in oocytes in Prophase I. **(a)** Angle evaluation and interpretation. The angle is computed as the angle between the droplet displacement and the vector of inward radial direction. When angles are comprised between 90 and 270° (red), the movement is toward the cell center. When angles are comprised between 0 and 90° or 270° and 360° (green), the movement is toward the cell cortex. **(b)** Distribution of ∆r.cos(θ) and ∆r.sin(θ) for droplets injected in oocytes maintained in Prophase I and treated or not with Cytochalasin D and observed at short time scale (∆t = 500 ms). Angles were evaluated at 1 ∆t. n = 29 oocytes in Prophase I and n = 14 oocytes in Prophase I + Cytochalasin D. The gray dashed line represents the zero of the distribution. **(c)** Bias toward the center is detected only in presence of F-actin (green). Velocity and diffusion coefficient can be computed. The nucleus is in blue, the oil droplet in yellow. **(d)** Quantification of the Mean Square Displacement (MSD in µm^2^) of the droplet centroid as a function of the delay (in sec) for droplets injected in Prophase I and observed at short time scale. n = 29 oocytes in Prophase I and n = 14 oocytes in Prophase I + Cytochalasin D. Mean (dark black for Prophase I and dark blue for Prophase I + Cytochalasin D curves) and standard deviation (grey and light blue quadrants) are presented. **(e)** Oil droplet moving toward the center in a Prophase I oocyte (observation at long time scale, transmitted light images, ∆t = 20 minutes). Images correspond to the Supplementary Video 3. One frame is shown every 100 min. Scale bar is 10 µm. **(f)** Trajectories of centroids from droplets centered during the experiment. Time (in min) is encoded in color on the centroid trajectories. The coordinates of the droplet centroid are given in the oocyte referential where (0,0) is the oocyte center. n = 22 oocytes. **(g)** Distance of the droplet centroid to the oocyte center presented as a function of time (in min) for each individual droplet (grey curves). The black line represents the mean distance to the oocyte center as a function of time (in min) for all droplet centroid trajectories. n = 22 oocytes.

From the mean value of the radial displacement ∆r.cos(θ) over a time step Δt, we obtained a mean instantaneous velocity in the presence of F-actin of 0.085 μm.min^-1^. In turn, from the standard deviation of either the radial or orthoradial components of the displacement, we could evaluate the diffusion coefficient of the droplets defined as 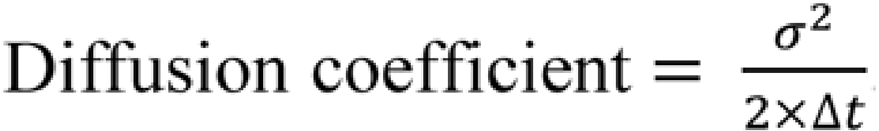. For droplets injected in oocytes in Prophase I, we found a diffusion coefficient of 0.071 µm^2^.min^-1^, value consistent with the one obtained from a linear fit of the Mean Squared Displacement (hereafter called MSD, Fig. 1d) of the oil droplets observed at longer time scales (0.5 to 200 s). Indeed, in the presence of F-actin, the MSD displays a linear dependence on time (Supplementary Fig. S1c and S1d), indicating a diffusive random motion (Fig. 1d), with a fitted diffusion coefficient of 0.072 µm^2^.min^-1^ (Supplementary Fig. S1c and e). On the contrary, in oocytes treated with Cytochalasin D, the droplets are almost immobile, with an estimated diffusion coefficient 10 times lower than controls (Fig. 1d compare black and blue curves; Supplementary Fig. S1c and Supplementary Video 2). This shows that the motion of oil droplets inside the cytoplasm depends on F-actin. Since the dynamics of the cytoplasmic actin mesh is controlled by the Myosin Vb motor^19^, this observation confirms that oil droplets undergo active (i.e. non thermal) diffusion, as observed for vesicles^2^. In conclusion, oil droplet movement can be described by a biased diffusion towards the oocyte center, which is mediated by the activity of the F-actin meshwork (Fig. 1c).

To further describe the ultimate displacement of oil droplets, we observed them in Prophase I oocytes at longer time scales (15 hours) and with a lower temporal resolution (20 minutes). Interestingly, oil droplets were progressively centered (Fig. 1e, Supplementary Video 3), and remained in the proximity of the oocyte center (Supplementary Fig. S2). The oil droplet injection technique allows following a wide range of droplet sizes. In our case, we managed to produce droplets with diameters comprised between 5 to 35 µm (Supplementary Fig. S3), comparable in size to the nucleus diameter (25 µm from ^2^). 76% of the droplets were centered during the 15 hours-duration of the experiment (Fig. 1f and 1g, see Methods). This is similar to the rate of nucleus centering in *Fmn2^-/-^* oocytes injected with Formin 2; indeed, 74% of the nuclei reached the center during a similar observation time^2^. Additionally, even if droplets do not reach the center, all show a biased movement toward the central area of the oocyte, as observed for nucleus centering^2^. Together, our results demonstrate the general nature of the F-actin dependent centering mechanism in mouse oocytes in Prophase I.

### The oil droplet recapitulates the nucleus movement

When oil droplets were injected in the presence of an F-actin probe (GFP-UtrCH^21^), we observed bright actin filaments around the droplet and around the nucleus. The meshwork around the droplet seemed as dense as the meshwork around the nucleus, and was dependent on the level of expression of the F-actin probe (Fig. 2a; high expression levels on the left panels and moderate expression levels on the right panels). To quantify the increase in density of the actin meshwork, we measured intensity profiles on the droplet or on the nucleus, in condition of high levels of GFP-UtrCH expression, and compared to the intensity in the cytoplasm (Methods). This analysis confirmed a quantitatively comparable enrichment of F-actin around both the nucleus and the droplet (Fig. 2b). This enrichment of F-actin is about 50% larger around the droplet or nucleus as compared to the bulk of the cytoplasm (Fig. 2b).

**Figure 2.**
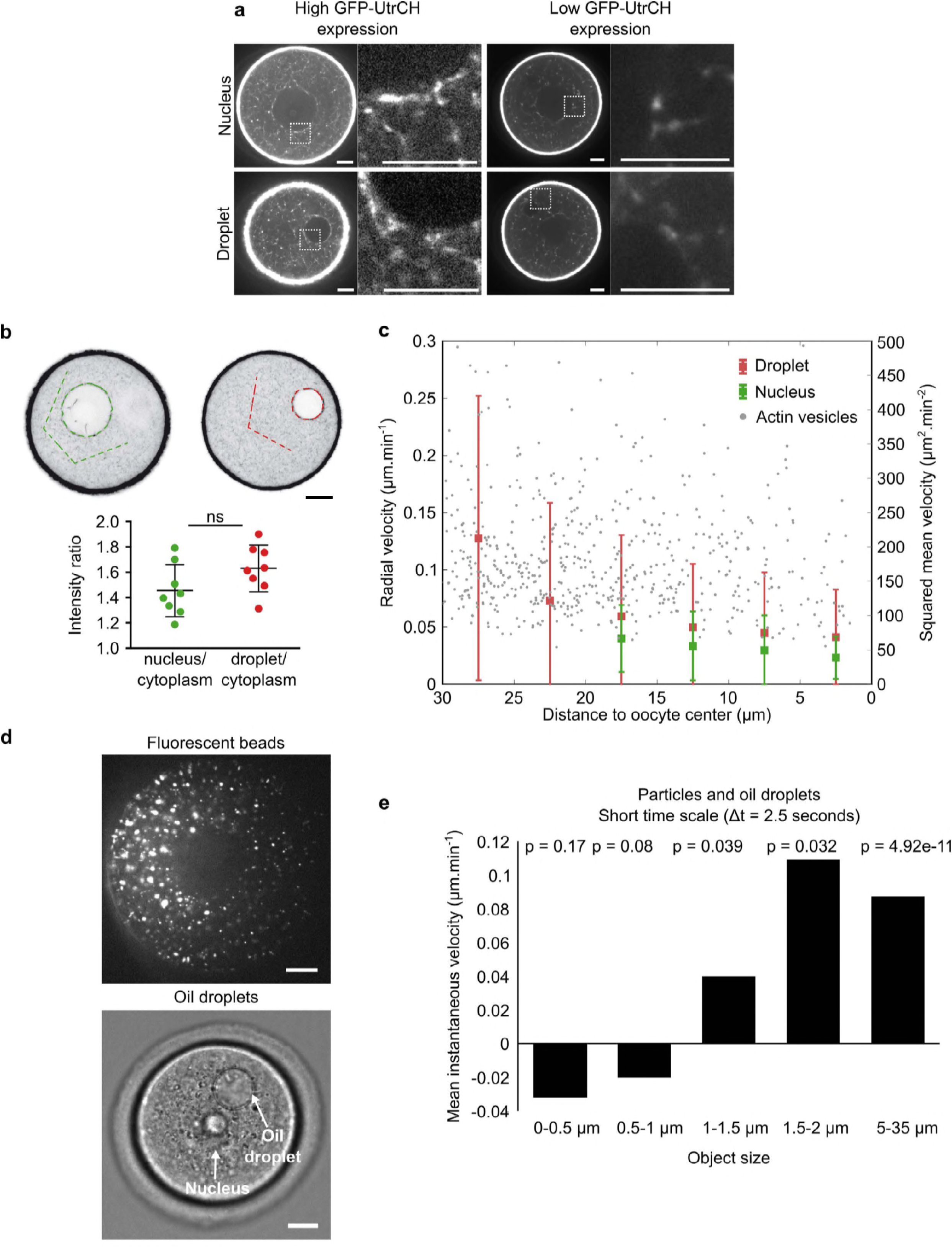
Oil droplets recapitulate nucleus behavior. **(a)** Visualization of F-actin in the region of the nucleus (upper panels) and of the droplet (lower panels) observed in the same oocyte in two different Z planes. F-actin is labelled with GFP-UtrCH, which is either highly-expressed (left panels) or midly-expressed (right panels). The square dashed region is enlarged on the right panels for each condition of GFP-UtrCH expression. F-actin is in white. Scale bars are 10 µm. **(b)** Quantification of the GFP-UtrCH fluorescence intensity ratio between the nucleus and the cytoplasm (left) compared to the one between the droplet and the cytoplasm (right). T-test resulted in a non-significant difference between the two distributions (p= 0.78). F-actin is in black. Scale bar is 10 µm. **(c)** Red (left y-axis): Radial velocity of oil droplet centroid for droplets with a diameter between 20 and 30 µm. n = 6 oocytes. Green (left y-axis): Radial velocity of nucleus centroid for Fmn2^-/-^ oocytes injected with Formin 2. Mean and standard deviation are superimposed. Grey (right y-axis): squared instantaneous velocity of actin vesicles (data from ^2^). **(d)** Top: image of fluorescent beads aggregated in a Prophase I oocyte. They correspond to objects with a diameter between 100 nm and 2 µm. Bottom: image of an oil droplet injected into a Prophase I oocyte. They correspond to objects between 5 and 35 µm in diameter. The oocyte nucleus is 25 µm wide. Scale bars are 10 µm. **(e)** Mean instantaneous velocity as a function of the object size. Velocity is computed from the distribution of ∆r.cos(θ) at 5 ∆t (2.5 s). P represents the p-value, probability that the distribution is significantly different from a normal distribution with the same standard deviation and centered at 0 (result of a z-test). For objects below 1 µm, there is no bias toward the center. For objects above 1 µm, there is a significative bias toward the oocyte center. n = 5 oocytes for aggregates of particles, n = 29 oocytes for oil droplets.

We then compared the radial velocities (defined as the radial component of the displacement vector over elapsed time) of droplets having a diameter between 20 and 30 µm, in the range of the nucleus diameter (Fig. 2c), to the radial velocities obtained for the nucleus velocity in Formin 2 knockout (*Fmn2^-/-^*) oocytes re-expressing Formin 2^2^. First, the radial velocities for droplets computed at long time scales (5-10 hours) were found in the range 0.04 and 0.13 µm.min^-1^ (Fig. 2c), and therefore of the same order of magnitude as the mean instantaneous velocity obtained for Δt = 500 ms (0.085 µm.min^-1^; Supplementary Fig. 1a). We observed that the radial velocities were slightly higher for oil droplets than for the nucleus (Fig. 2c, compare green and red points). In both cases we observed a decrease of velocity at the center, consistent with a vanishing bias. The radial velocities for droplets have a larger variability than the one measured for the nucleus. These slight quantitative differences could be due to different effects. First, the diameter of oil droplets is more variable than the nucleus diameter, since not genetically encoded but controlled manually (see Methods). Second, in the case of the nuclear repositioning, cRNA encoding for Formin 2 was injected into *Fmn2^-/-^* oocytes and thus nucleus centering is occurring while the F-actin mesh is progressively reforming. In contrast, in the case of oil droplets, these are injected at steady state in control oocytes that already present a fully dynamic actin mesh. Third, the droplet is made of incompressible oil and has different mechanical properties as compared to a nucleus. Fourth, we cannot exclude that the composition of the nuclear membrane itself has an own contribution, not recapitulated in our assays. Nevertheless, our results suggest that oil droplets display a dynamic behavior very close to that of the nucleus.

We then wanted to test the impact of the size of objects on their centering efficiency. We studied objects with different ranges of size. First, we used the oil droplets to produce large objects (between 5 to 35 µm in diameter). Second, we used 100 nm fluorescent particles; when injected at a high concentration (Methods), they aggregate in the cytoplasm of the oocyte (Fig. 2d). We took advantage of this aggregation allowing the production of objects with sizes between 100 nm to 2 µm (measured by the fluorescence intensity of the detected spot; see Methods), the injection of very large particles being technically challenging.

We computed the MSD for each individual object and extracted the diffusion coefficient from a linear fit over a 0-10 s range (Supplementary Fig. S4b). The obtained values of diffusion coefficient were consistent with those deduced from the variance of displacement vectors at 1 ∆t. Interestingly, for small objects, the diffusion coefficient does not depend on the size of the object. This is in agreement with what was found previously for cellular clusters in *Drosophila*^22^ embryos.

For each trajectory, we analyzed the radial and orthoradial component of the object displacement. From the distribution, we computed the mean instantaneous velocity for a delay of 2.5 s (5 ∆t), and found that it increases with the object size (Fig. 2e). Comparing the distributions of radial components to Gaussian distributions centered at 0, revealed that there is a significant bias for objects presenting a diameter larger than 1 µm. Altogether, the results of the bias and the angle analysis argue that there is a cut-off size of 1 µm for the centering mechanism at short time scales. Notably, this is consistent with previous observations on much longer time scales (18 hours period), which also suggested that objects smaller than 1 µm do not center^2^. This supports the existence of a non-specific centering mechanism, experienced by objects larger than 1 µm.

### Oil droplets are centered in oocytes that undergo Meiosis I

As introduced previously, a question unresolved in the field is how the same molecules (Formin 2, Spire 1 & 2 and Myosin Vb) promote two opposite motions: centering of chromosomes in Prophase I^2^ and off-centering of chromosomes later in Meiosis I^7,10,16–19^. We thus decided to address whether the centering mechanism was maintained during Meiosis I. To this aim, we injected oil droplets in Prophase I oocytes, then allowed oocytes to synchronously resume meiosis (evidenced by Nuclear Envelope BreakDown, NEBD) and to proceed into Meiosis I. Meiosis I is a long process taking about 10h and is described in min after NEBD (Fig. 3a). First, we observed that oocytes could undergo Meiosis I unperturbed, since they succeeded in dividing on time and extrude a first polar body (PBE), arguing that the oil droplet is not toxic for the development of the oocyte (Fig. 3a). It also showed that the oil droplet could co-exist in the cytoplasm with the mechanism that promotes off-centering of the first meiotic spindle, a pre-requisite for the first asymmetric division, i.e. first polar body extrusion^5^. Second, we observed that the oil droplet is being centered during the process of Meiosis I (Fig. 3a, Supplementary Video 4). Effectively, 92% (11 out of 12) of the droplets were centered before the extrusion of the first polar body (Fig. 3c and 3e). When the actin network is dismantled (in the presence of Cytochalasin D) no movement of the droplet was observed, arguing that droplet centering is a consequence of the presence of actin (Fig. 3b, 3d, 3f, Supplementary Video 5). This phenomenon is unexpected because it means that during Meiosis I, when the spindle is migrating towards the cortex, a centering mechanism is nonetheless present. Surprisingly, the same molecular actors are in action as Myosin Vb was shown to play a key role in the nucleus centering mechanism^2^ and in spindle migration^19^. These results show the co-existence of two mechanisms in Meiosis I: a specific one that allows off-centering of the spindle toward the cortex, involving Myosin II activity^16–18^ and a non-specific one that ensures the centering of big objects on long time scales. To have a better understanding of this process, we compared the characteristics of droplet centering in Prophase I and in Meiosis I.

**Figure 3.**
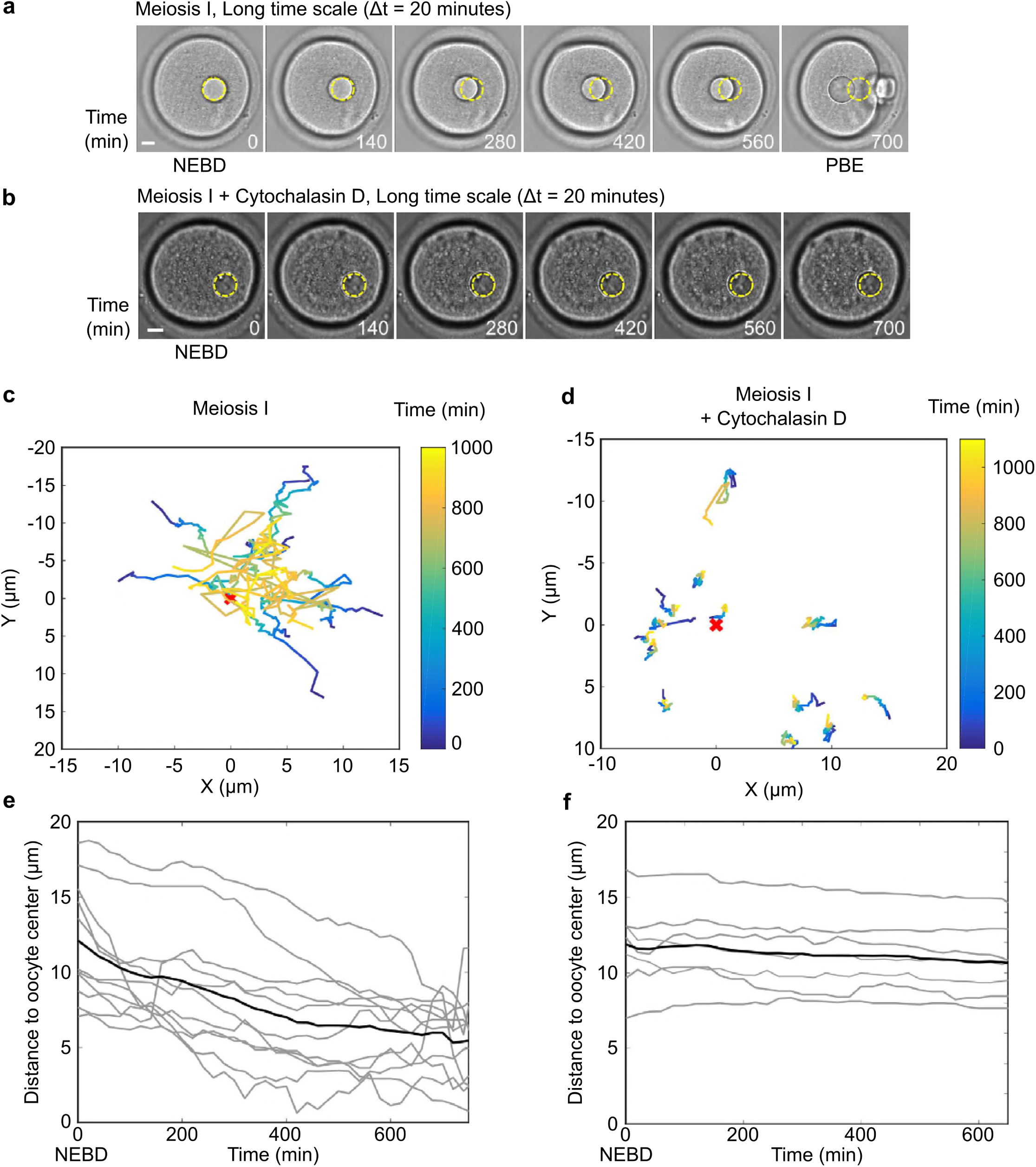
Oil droplets are also centered in oocytes undergoing Meiosis I. **(a)** Centering of an oil droplet in an oocyte undergoing Meiosis I. The images correspond to Supplementary Video 4. One frame is shown every 140 min. The first frame corresponds to NEBD (Nuclear Envelope BreakDown), a marker of meiosis resumption and the last one to PBE (Polar Body Extrusion), a marker of meiosis I completion. The initial location of the oil droplet is highlighted by a dotted-yellow circle on each picture. Scale bar is 10 µm. **(b)** An oil droplet observed during Meiosis I in an oocyte treated with Cytochalasin D. The images correspond to Supplementary Video 5. One frame is shown every 140 min. The first frame corresponds to NEBD. Note that oocytes treated with CCD do not extrude a polar body (PBE). The initial location of the oil droplet is highlighted by a dotted-yellow circle on each picture. Scale bar is 10 µm. **(c)** Trajectories of droplet centroids that are centered during the observation in Meiosis I. Time (in min) is encoded in color on the centroid trajectories. The coordinates of the centroid droplet are given in the oocyte referential where (0,0) is the center of the oocyte. n = 11 oocytes. **(d)** Trajectories of droplet centroids, not centrally located at the beginning of the observation, in Meiosis I oocytes treated with Cytochalasin D. Time (in min) is encoded in color on the centroid trajectories. The coordinates of the centroid droplet are given in the oocyte referential where (0,0) is the center of the oocyte. n = 13 oocytes. **(e)** Distance of the droplet centroid to the oocyte center presented as a function of time (in min) for each individual droplet for oocytes in Meiosis I (grey curves). The black line represents the mean distance to the oocyte center as a function of time (in min) for all droplet centroid trajectories. n = 11 oocytes. **(f)** Distance of the droplet centroid to the oocyte center presented as a function of time (in min) for each individual droplet for oocytes in Meiosis I treated with Cytochalasin D (grey curves). The black line represents the mean distance to the oocyte center as a function of time (in min) for all droplet centroid trajectories. n = 13 oocytes.

### Comparison of droplet centering in Prophase I versus Meiosis I

To compare the process of droplet centering between the two stages, we first analyzed the radial velocity for droplets observed in Prophase I or in Meiosis I (Fig. 4a). Droplets injected in oocytes undergoing Meiosis I show a slower radial velocity than the droplets injected in oocytes in Prophase I. To have a quantitative characterization of the centering process, we computed a mean centering velocity (Methods; Fig. 4b upper panel) with t = 0 corresponding to the beginning of the movie for Prophase I and t = 0 corresponding to NEBD for Meiosis I. We observed that the mean centering velocity was 4 times smaller in Meiosis I than in Prophase I (0.10 µm.min^-1^ in Prophase I compared to 0.024 µm.min^-1^ in Meiosis I, Fig. 4b lower panel). The higher dispersion of the data in Prophase I could be explained by the larger diameter range of droplets injected in Prophase I versus Meiosis I (Supplementary Fig. S5). Therefore, even if droplets were centered in both cases, the characteristics of centering are not exactly similar. In Prophase I, the centering displays more noise but occurs faster. The differences in centering could be due to the fact that in Prophase I we perform experiments at steady state while in Meiosis I the mesh is progressively reforming while droplets are followed^8^.

**Figure 4.**
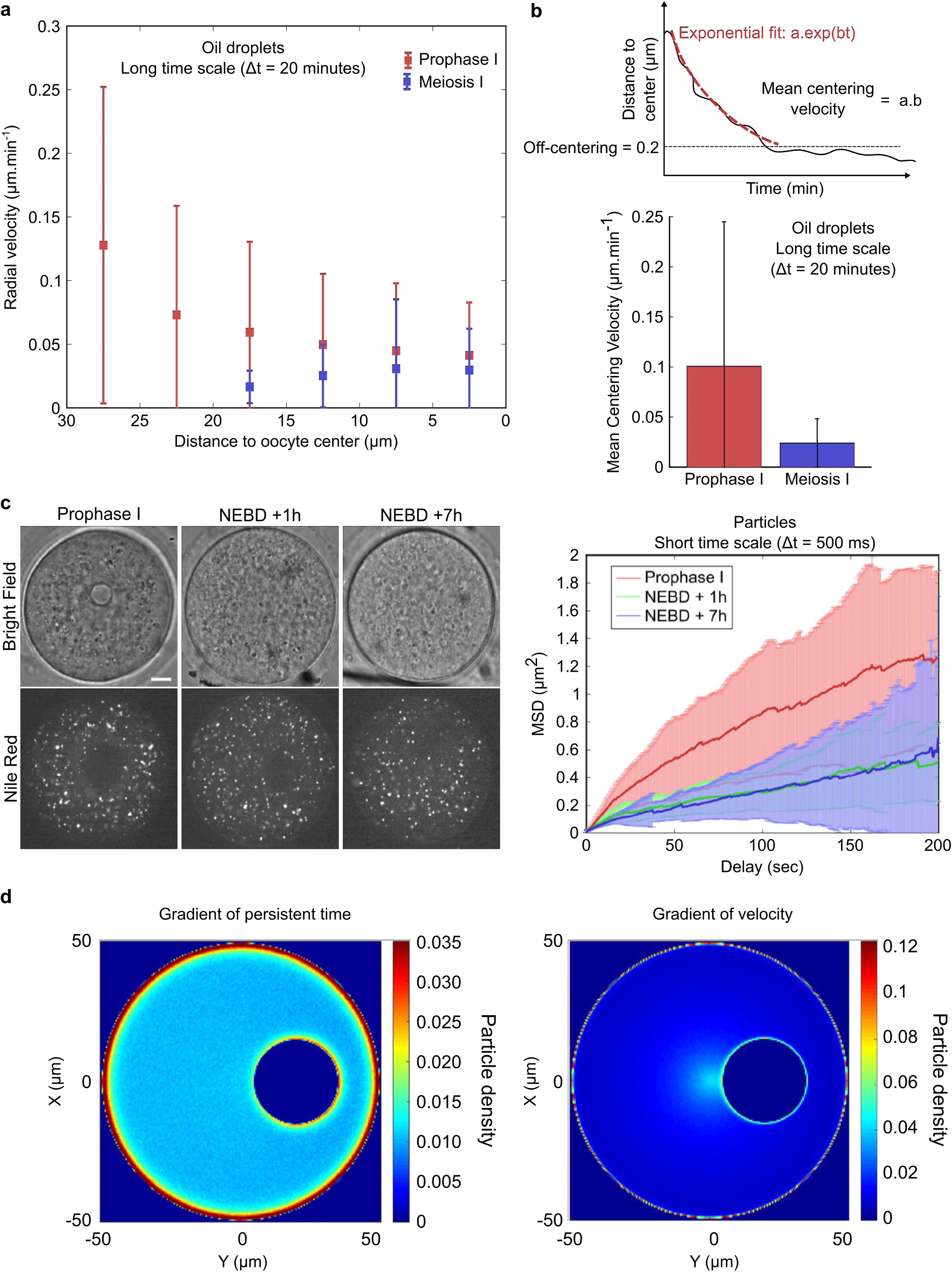
Comparison of droplets centering in Prophase I versus Meiosis I. **(a)** Radial velocities of droplet centroids as a function of distance to oocyte center for droplets centered during Prophase I (red dots) and Meiosis I (blue dots). Mean and standard deviation are represented. n = 22 oocytes in Prophase I and n = 11 oocytes in Meiosis I. **(b)** Top: Scheme explaining the computation of the centering velocity by an exponential fit on the distance to center as a function of time. Bottom: average of the centering velocity (in µm. min^-1^) for droplets centered during the experiment in Prophase I (red histogram) and Meiosis I (blue histogram on the Bottom panel). n = 22 oocytes for Prophase I and n = 11 oocytes for Meiosis I; error bars show standard deviation. **(c)** On the left pictures, fluorescent labelling of total vesicles using Nile Red, in oocytes observed in Prophase I, at early (NEBD + 1h) or late (NEBD + 7h) Meiosis I. Bright field is presented on the upper row and fluorescent images on the lower row. Scale bar is 10 µm. On the right panel, MSD (in µm^2^) of the total vesicles for the various conditions of the left panel is measured as a function of the delay (in sec). Mean (dark red for Prophase I, dark green for NEBD +1h and dark blue for NEBD +7h curves) and standard deviation (light red for Prophase I, light green for NEBD +1h and light blue for NEBD +7h quadrants) are presented. **(d)** Comparing activity gradient due to either a persistence time or a velocity. Scheme of theoretical prediction for active particle density and accumulation around objects, for two limit cases. The average particle density from a simulation of a circular 2D system of active Brownian particles with a circular object inside it is plotted for: (left) a uniform speed υ, and a persistence time τ linearly increasing from the center to the cortex, (right) a uniform τ, and υ linearly increasing from the center to the cortex. On the left, the bulk density is uniform while the accumulation of particles around the disk is not. On the right, the bulk density is inversely proportional to the local υ, while the edge accumulation of particles is uniform due to the uniform τ. The heat map represents the particle density (number of particles per unit area) in both the left and the right panel. The range of the heat map was selected in each panel so that the particle accumulation around the disk is clearly visible, namely so that it is clear that the particle density around the disk is uniform in the υ gradient case but non-uniform in the τ gradient case. (Simulations were performed as described in ^13^). Simulation parameters: *R* = 50, *R*_*d*_ = 15, *r*_*d*_ = 20, *γ* = 10, *D* = 1, *N* = 100, simulation step size *dt* = 1*e* − 4, total runtime 2.5e6. 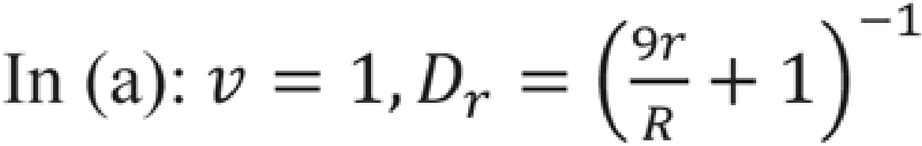, 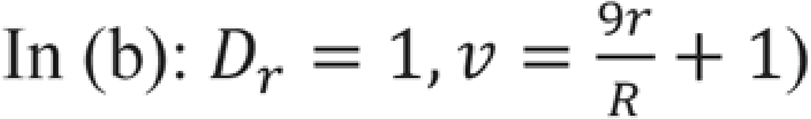

We next checked if we were able to link those data with the rheological properties of the cytoplasm. We first performed active micro-rheology experiments using optical tweezers on endogenous vesicles to probe the mechanical response of the oocyte cytoplasm in Prophase I and Meiosis I (Supplementary Fig. S6). We found no significant difference in the elastic modulus (G’) between oocytes observed in Prophase I and Meiosis I (Supplementary Fig. S6, left panel). As for the viscous modulus (G’’), a slight difference between Prophase I and Meiosis I oocytes was observed (slightly higher in Prophase I, Supplementary Fig. S6, right panel). Note that we obtained a viscosity of the order of 1 Pa. s for the cytoplasm of oocytes maintained in Prophase I, similar to what was previously found with magnetic tweezers measurements^23^. This analysis suggested that oocytes in Prophase I or Meiosis I have comparable mechanical responses, which alone cannot explain the observed difference in the centering speed.

To understand the differences of behavior between Prophase I and Meiosis I and probe the spontaneous dynamics of the cytoplasm, we followed the movement of endogenous vesicles labeled with Nile Red, a fluorescent dye (Fig. 4c and Supplementary Video 6). We computed the MSD and extracted the effective diffusion coefficients for Nile Red positive vesicles (Fig. 4c, and Supplementary Fig. S7). In Prophase I, vesicles have a diffusion coefficient of 0.23 µm².min^-1,^ while in Meiosis I, they display a diffusion coefficient of 0.12 µm².min^-1^ at NEBD+1h (actin meshwork dismantled) and 0.09 µm².min^-1^ at NEBD +7h (actin meshwork reassembled), indicating an increased activity in Prophase I. This finding, together with the observation of a (4 times) faster mean centering velocity in Prophase I (Fig. 4b, c) supports our hypothesis that the centering mechanism is controlled by cytoplasmic actin activity. Indeed, as the optical tweezer experiments show that the mechanical response of the cytoplasm is approximately unchanged (Supplementary Fig. S6), the observed changes in effective diffusion can be attributed to active forces on the vesicles. Hence the activity of the total vesicles, and therefore of the cytoplasm, is found to be correlated with the centering velocity of a large object like a nucleus or an oil droplet: in the case of lower cytoplasmic activity, the centering is slower, as observed for Meiosis I (Fig. 4b).

### Comparison to theoretical analysis in terms of active Brownian particles

Altogether, our observations are qualitatively consistent with the physical description introduced in ^2,12,13^, where the non-uniform cytoplasmic activity is quantified by a gradient of velocity of actin-positive vesicles. In these theoretical descriptions, the observed gradient of activity leads to a non-specific effective pressure force toward the oocyte center acting on any particle immersed in the cytoplasm. This active pressure, can be treated in a simplified manner as a collection of self-propelled particles (actin vesicles) that have a distinct propulsion force (giving them an intrinsic velocity υ) and persistence time τ, which are a function of the distance from the cell center. Such a simple microscopic model for the spatially inhomogeneous diffusion of the actin vesicles can be analyzed by numerical simulations (Fig. 4d) and can hint at the origin of the inhomogeneity and provide predictions that can be tested experimentally^13^.

Within such models, since the persistence length *l*_*p*_ = *vτ* of the vesicle motion is small compared to other length scales in the system^13^, the force with which the vesicles pushes an object, such as the nucleus or a droplet, is predicted to be proportional to the gradient in persistence length ∇*l*_*p*_, directed towards decreasing values of *l*_*p*_. The two simple limits to achieve a gradient in *l*_*p*_, are to have a gradient in either one of *v* and τ, while the other is held constant^13^. In the oocyte, both parameters could in principle vary spatially.

Because of the persistence of their motion, active particles tend to accumulate on boundaries and surfaces. The accumulation is proportional to the local persistence time τ, and therefore, if there is a gradient in τ the accumulation of particles around a circular object should be larger in the larger τ regions (as in Fig. 4d left). In contrast, if there is a gradient in *v* the accumulation should be approximately uniform around the object (Fig. 4d right). The approximately uniform vesicle density (shown in Supplementary Fig. S4a of ^2^) and the existence of a force pushing objects towards the center are consistent with a uniform *v* and a radially varying τ. Actin accumulation around the droplets was observed (Fig. 2a), however from the available data it could not be determined if there is a bias in actin accumulation. We could sometimes observe around the droplet a slight bias with more actin facing the cortex than the oocyte center, but not in a consistent manner (two examples presented in Supplementary Fig. S4a). Future work using better resolved imaging conditions could test this prediction.

Alternatively, a system with a radial gradient in υ could have a roughly uniform density of vesicles (as observed^2^) only if there are rapid processes of vesicle formation and annihilation throughout the oocyte cytoplasm^12^. The currently available data about the life-time of the vesicles is not precise enough to rule this out, and this scenario would predict that the accumulation of actin on the surface of the nucleus (or droplet) has no center-cortex bias. Future studies with high temporal resolution of the vesicles’ trajectories in three dimensions could allow to determine their life-time, and test the υ-gradient scenario. Comparing the results of these models to experiments will then help direct future studies of the mechanism used by the cell to produce the gradient in activity, which remains an open puzzle. Simulations of the pressure exerted by a gas of self-propelled particles, with a gradient in either the intrinsic velocity or the persistence time, provide detailed predictions potentially revealing the mechanism for the activity gradient in the oocyte.

## Discussion

By using oil droplets that mimic the nucleus (size, with F-actin recruitment), we demonstrate that the centering mechanism is non-specific to the biological nature of the nucleus. In Prophase I, oil droplets and nucleus centering have the same characteristic (time, velocity etc.). We also show that the centering mechanism is still present in Meiosis I, with slower kinetics than in Prophase I.

We tested the influence of object size on the centering efficiency. Indeed, large objects (above 1 µm) show a biased movement toward the oocyte center at short time scale (2.5 sec). Interestingly this cut-off is of the same order of magnitude as the estimated meshwork size for oocytes maintained in in Prophase I^8,16^. At short time scale (sec), large objects show a more restricted displacement compared to small objects, probably due to a friction effect. At long time scale (hours), for oil droplets (objects between 5 and 35 µm in diameter), the centering velocity does not depend on the oil droplet size (Supplementary Fig. S5). This suggests that friction forces scale much faster than linearly with particle size, as has been observed in *Drosophila* embryos^22^, and in contrast to the behavior expected in the low-Reynolds Stokes regime.

The effectiveness of the centering mechanism can be quantified by the Peclet number, which quantifies whether the bias toward the center is stronger than diffusion for our objects. Peclet number is given by 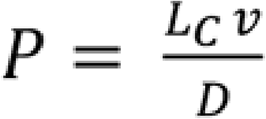 with *v* the velocity of the object, *Lc* the characteristic length of the system (20 µm here), and *D* the diffusion coefficient. *v* can be estimated from the long time scale movies (to ensure that we measure a biased velocity and not a fluctuation velocity). In Prophase I, we found *v* = 0.060 µm.min^-1^. For oil droplets in Prophase I, the diffusion coefficient (0.072 µm^2^.min^-1^) was estimated from the MSD obtained on short time scale movies (Supplementary Fig. S1). Therefore, Peclet number can be estimated at *P* ≈ 17 and thus we can conclude that for oil droplets, the bias is dominating the fluctuations. Our work provides another example of how F-actin acts as an organizer of the intracellular space, by distributing objects in space as a function of their size. For example in the *Xenopus* oocyte nucleus, F-actin has been shown to act as a stabilizing scaffold that would prevent the influence of gravity^24^.

The slower droplet centering in Meiosis I could be explained by the fact that at this stage, the oocyte shares actin resources for two processes: a non-specific centering mechanism and a specific off-centering of the meiotic spindle via pulling forces exerted by Myosin II at spindle poles and through specific F-actin connections between the meiotic spindle poles and the cortex^16–18^. Sharing of resources for actin networks has been widely studied in fission yeast^25,26^. It is also interesting to consider the centering mechanism during Meiosis I as a mechanism that counteracts spindle migration. Indeed, if spindle migration was dragging all the maternal stores into the polar body, then the asymmetric division would deplete the oocyte from the reserves necessary for future embryo development. It is possible that the centering mechanism in Meiosis I is a safe-guarding mechanism to preserve most organelles and RNPs granules in the oocyte itself instead of being transported into the polar body.

## Methods

### Oocyte collection, culture and microinjection

Oocytes were collected from 11 week-old OF1 mice as previously described^5^ and maintained in Prophase I arrest in M2+BSA medium supplemented with 1 μM Milrinone^27^. All live culture and imaging were carried out under oil at 37°C. We used the following pspe3-GFP-UtrCH^7^ construct to produce cRNA. *In vitro* synthesis of capped cRNAs was performed as previously described^5^. cRNAs were centrifuged at 4°C during 45 minutes at 13000 rpm before microinjection. The cRNA encoding GFP-UtrCH were injected first and then oocytes were injected with oil droplets. We injected Fluorinert FC-70 (Sigma, Ref. F9880) with a density of 1.9 times the density of water at the maximal pressure of the microinjector (Clean mode at 7000 hPa). The size of the oil droplets was visually adjusted by manual control of the duration of the microinjection pulse. Microinjections were performed using an Eppendorf Femtojet microinjector at 37°C as in^28^.

### Drug treatments

Cytochalasin D (Life Technologies, Ref. PHZ1063) was diluted at 10 mg.ml^-1^ in DMSO and stored at −20°C. It was used on oocytes at 1 μg.ml^-1^. Nile Red stain (Sigma, Ref. N3013) was used to label the total pool of vesicles. It was diluted at 5 mg.ml^-1^ in DMSO and stored at room temperature. It was used on oocytes at 10 μg.ml^-1^. For the injection at high concentration (to have aggregates formation) of latex fluorescent beads (0.1 µm; Life Technologies, F8803), the beads were rinsed several times in nuclease-free water before use to remove traces of sodium azide and diluted 10 times before injection.

### Live imaging

Spinning disk images were acquired at 37°C in M2 + BSA +1 µM Milrinone using a Plan-APO 40x/1.25 NA objective on a Leica DMI6000B microscope enclosed in a thermostatic chamber (Life Imaging Service) equipped with a CoolSnap HQ2/CCD-camera (Princeton Instruments) coupled to a Sutter filter wheel (Roper Scientific) and a Yokogawa CSU-X1-M1 spinning disk. For imaging oil droplets, we either acquired using the stream mode of the camera on Metamorph (one image every 500ms) or one image every 20 min to follow the whole motion towards the oocyte center. The actin cytoplasmic meshwork decorated with GFP-UtrCH and total vesicles stained with Nile Red were imaged every 500 ms with the stream acquisition mode of Metamorph upon excitation at 491 nm.

### Optical tweezer experiments

The single-beam gradient force optical trap system uses a near infrared fiber laser (λ= 1064 nm, YLM-1-1064-LP, IPG, Germany) that passes through a pair of acousto-optical modulators (AA-Optoelectronics, France) to control the intensity and deflection of the trapping beam. The laser is coupled into the beam path via dichroic mirrors (Thorlabs) and focused into the object plane by a water immersion objective (60x, 1.2 NA, Olympus). The condenser is replaced by a long distance water immersion objective (40x, 0.9 NA, Olympus) to collect the light and imaged by a 1:4 telescope on an InGaAs quadrant photodiode (QPD) (G6849, Hamamatsu). The resulting signal is amplified by a custom-built amplifier system (Oeffner Electronics, Germany) and digitized at a 500 kHz sampling rate and 16 bit using an analog input card (6353, National Instruments, Austin, TX, USA). The position of the trapped particle is measured by back focal plane interferometry^29^. All control of the experimental hardware is executed using LabVIEW (National Instruments). Optical trapping of endogenous (diameter ~ 1 micron) vesicles was calibrated using the active-passive method similarly as in^30^, where the high-frequency fluctuations (f > 300 Hz) are thermal in origin^31,32^. Vesicle size was estimated by comparing image analysis and laser interferometry profiles to 1 micron beads^32^. The mechanical response was measured by applying a sinusoidal force to a vesicle and observing the subsequent displacement. The shear modulus was calculated from the mechanical response using the generalized Stokes-Einstein relation as done previously^33^.

### Image Analysis

Image analysis was performed using ImageJ, Icy and Matlab. When needed, movies were realigned with the stackreg plugin of ImageJ. For all the automated tracking the software Icy was used^34^. Droplet tracking in bright field at short time scale was done with the Active Contours Plugin. Fluorescent particle and endogenous vesicle tracking were done with the Spot Detector plugin combined with the Spot Tracking and Track Manager plugins. The trajectories were exported in Excel files and analyzed with Matlab software. The tracks were filtered to keep only the tracks with more than 30 points (movies of 400 frames). This threshold was determined with simulated data, to avoid the tracking of false trajectories coming from the noise of the movie. Mean square displacement analysis was done with the msdanalyzer class in Matlab^35^. Diffusion coefficient was computed from a linear fit on the 20 first points of the MSD curve. The slope of the fit is equivalent to 4D with D the diffusion coefficient.

For the aggregated fluorescent particles, we computed the apparent diameter based on the intensity of the detected spot.

Estimation of centering: we defined that a droplet is centered when its off-centering is below 0.2, with the off-centering being the normalized distance compared to the oocyte radius (the off-centering is equal to 1 at the cortex and to 0 at the center of the oocyte). To compute the mean centering velocity, we fitted the distance as a function of time with an exponential function (*a*.exp(*b*.t)). The centering velocity is then defined as *v* = *a.b*.

The radial (∆rcos(θ)) and orthoradial (∆rsin(θ)) components were computed by taking the angle θ between the displacement (at a given ∆t) and the radial axis. ∆r is the norm of the displacement. Radial velocity is computed by taking the projection of the displacement on the radial vector over the time step.

To measure the intensity ratio between the droplet (or the nucleus) and the cytoplasm, we drew a contour line around the droplet (or the nucleus) and then a line of the same length in the cytoplasm. We then measured the intensity along these lines and computed the ratio between the droplet (or nucleus) intensity and the cytoplasmic intensity. For Fig. 2b, the t-test was performed with Prism. z-tests and Kolmogorov Smirnov statistical tests were performed with Matlab. For the interpretation of the p-values, NS means there is no significant difference between the two distributions. One star means p-value < 0.05, two stars means p-value < 0.01, and three stars means p-value <0.001.

## Acknowledgements

AC is supported by a “Ministère de la Recherche” doctoral fellowship. This work was supported by the ANR (ANR-DIVACEN to MHV & Renata Basto, Curie Institute, N°14-CE11), by an FRM Label (DEQ20150331758 to MHV), by a PSL “Aux Frontière des Labex” grant (MYOOCYTE, MHV as coordinator), by an Inca grant (PLBIO 2016-270-TRAN). This work has received support from the Fondation Bettencourt Schueller, support under the program «Investissements d’Avenir» launched by the French Government and implemented by the ANR, with the references: ANR-10-LABX-54 MEMO LIFE, ANR-11-IDEX-0001-02 PSL* Research University. NSG is the incumbent of the Lee and William Abramowitz Professorial Chair of Biophysics and acknowledges support from the ISF (Grant No. 580/12).

---

**Supplementary Figure S1.**
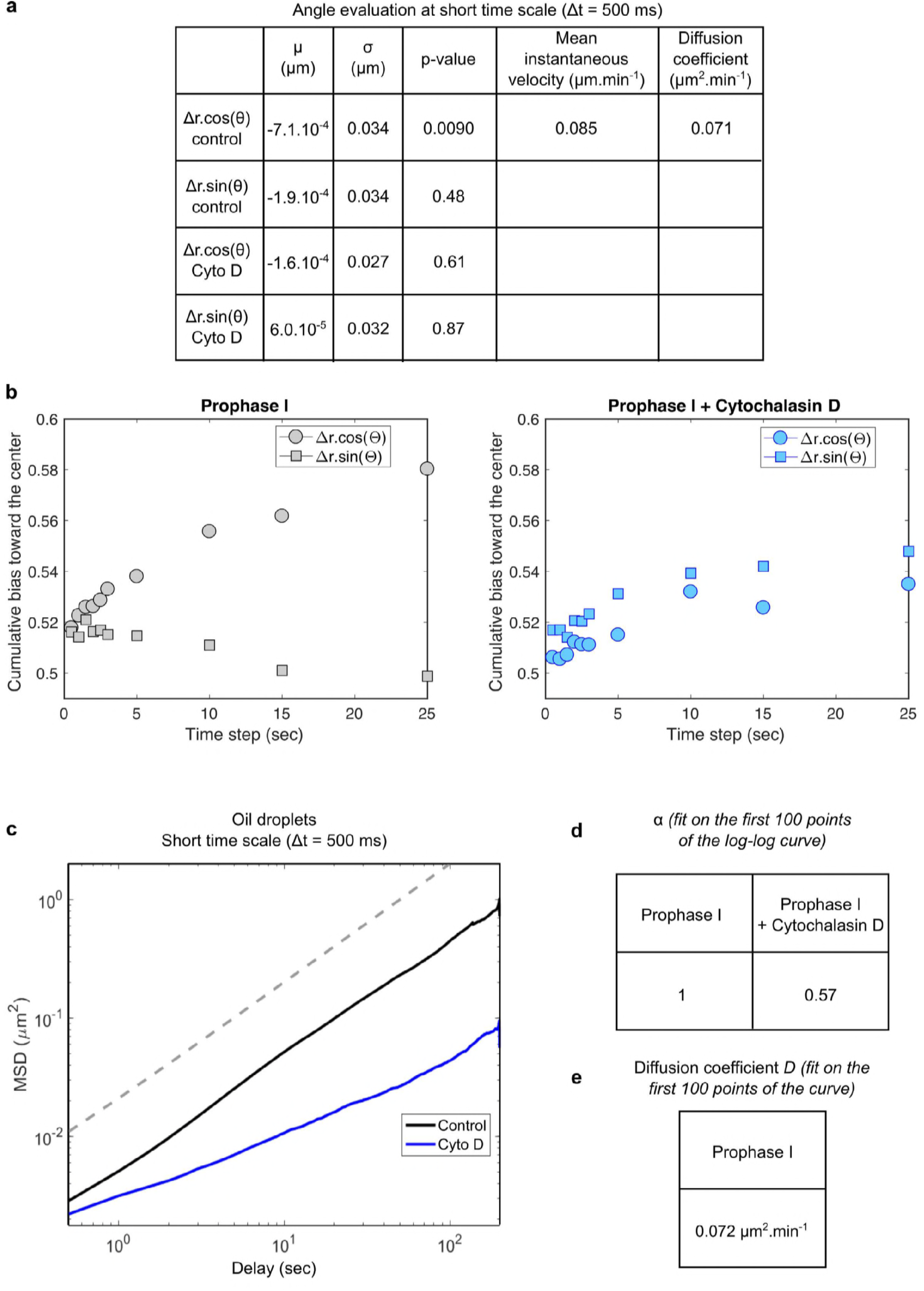
MSD interpretation for oil droplets observed at short time scales. **(a)** Estimation of parameters from the distributions of ∆r.cos(θ) and ∆r.sin(θ). µ represents the mean of the distribution, σ the standard deviation of the distribution. P-value is the probability that the distribution is significantly different from a normal distribution with the same standard deviation and centered at 0 (result of a z-test). The mean instantaneous velocity is computed as the mean of the distribution over the time (500 ms here). The diffusion coefficient is computed at the standard deviation to the power of two divided by the time by two.

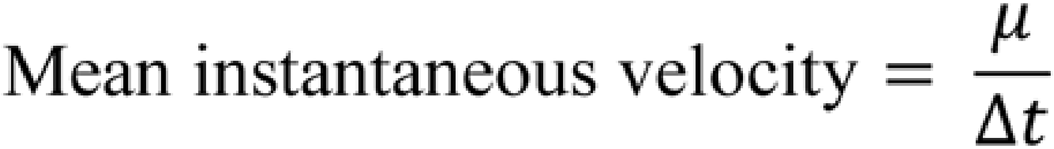

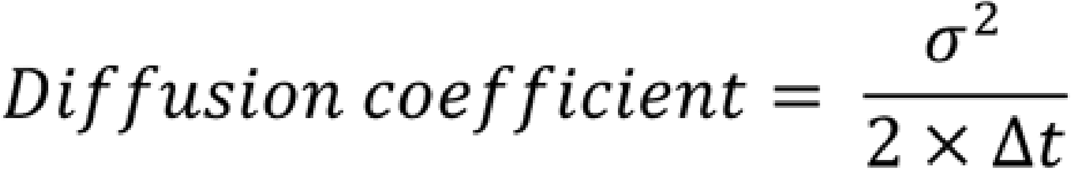 **(b)** Estimation of the cumulative bias toward the cell center (positive values here). Cumulative bias is computed for different delays and for radial and orthogonal components in oocytes maintained in Prophase I and in oocytes treated with Cytochalasin D. **(c)** Representation of the Mean Square Displacement (MSD) of oil droplets as a function of the delay (in sec) on a log-log scale in Prophase I at a short time scale. n = 29 oocytes in Prophase I (black curve) and n = 14 oocytes in Prophase I + Cytochalasin D (blue curve). The dotted line represents a linear regression with a slope of 1. **(d)** Results obtained from the fitting of the MSD on a log-log scale. α represents the slope of the curve (here the fit is made on the first 100 points of the curve). **(e)** Evaluation of the diffusion coefficient from the fit of the MSD as a function of time. The fit was done on the first 100 points of the curve. The diffusion coefficient is then evaluated considering the following formula: MSD = 4*D*t (with t the delay).

**Supplementary Figure S2.**
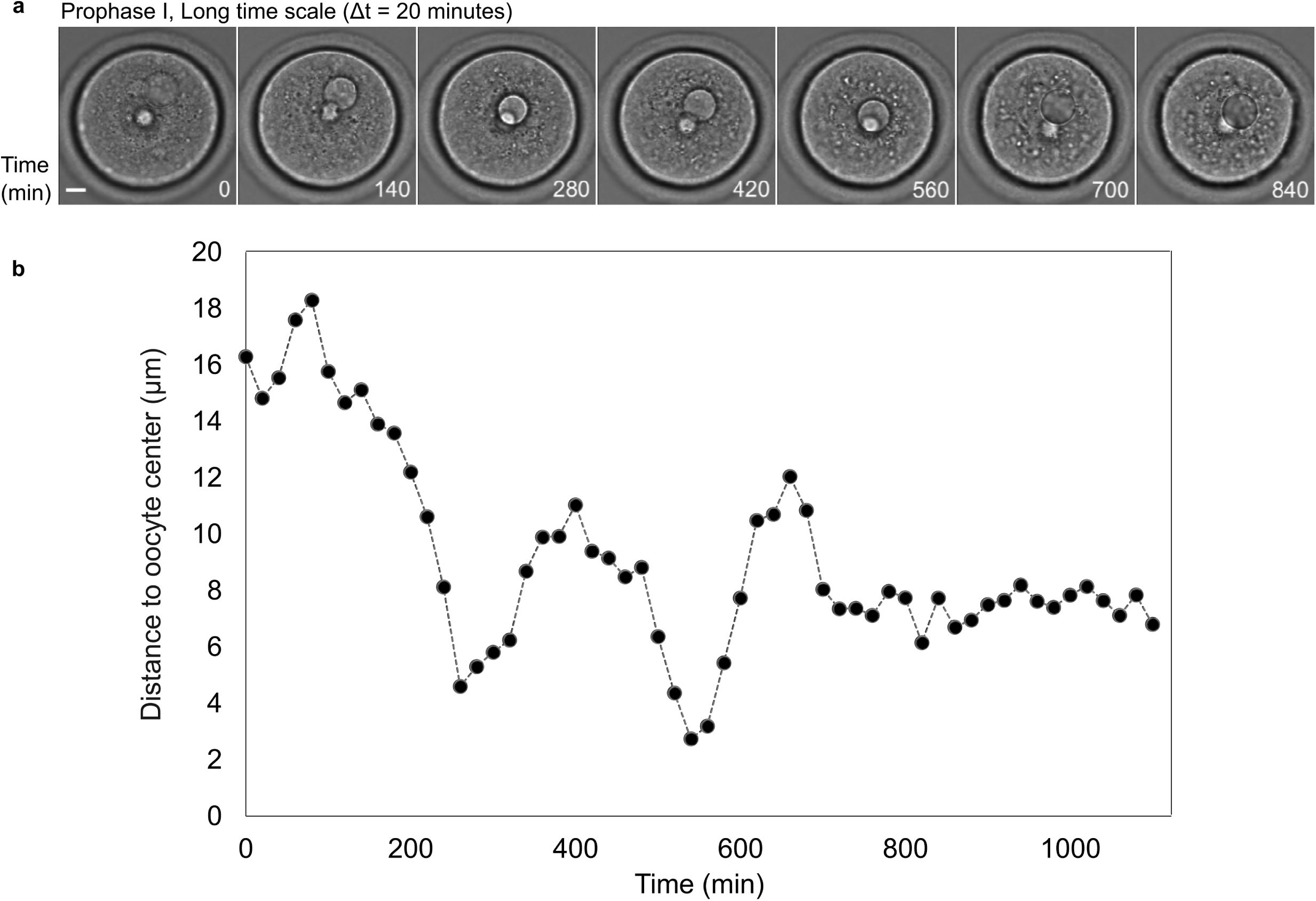
Example of an oscillatory motion of a droplet around the oocyte center. **(a)** Images in transmitted light of a droplet centered in Prophase I. One image is shown every 140 min. Scale bar is 10 µm. **(b)** Distance of the droplet centroid to the oocyte center as a function of time (in min) from the images shown in (a). Two oscillations of the droplet can be observed at 200 and at 600 min. It is worth noting that the droplet encounters the nucleus when it reaches the center.

**Supplementary Figure S3.**
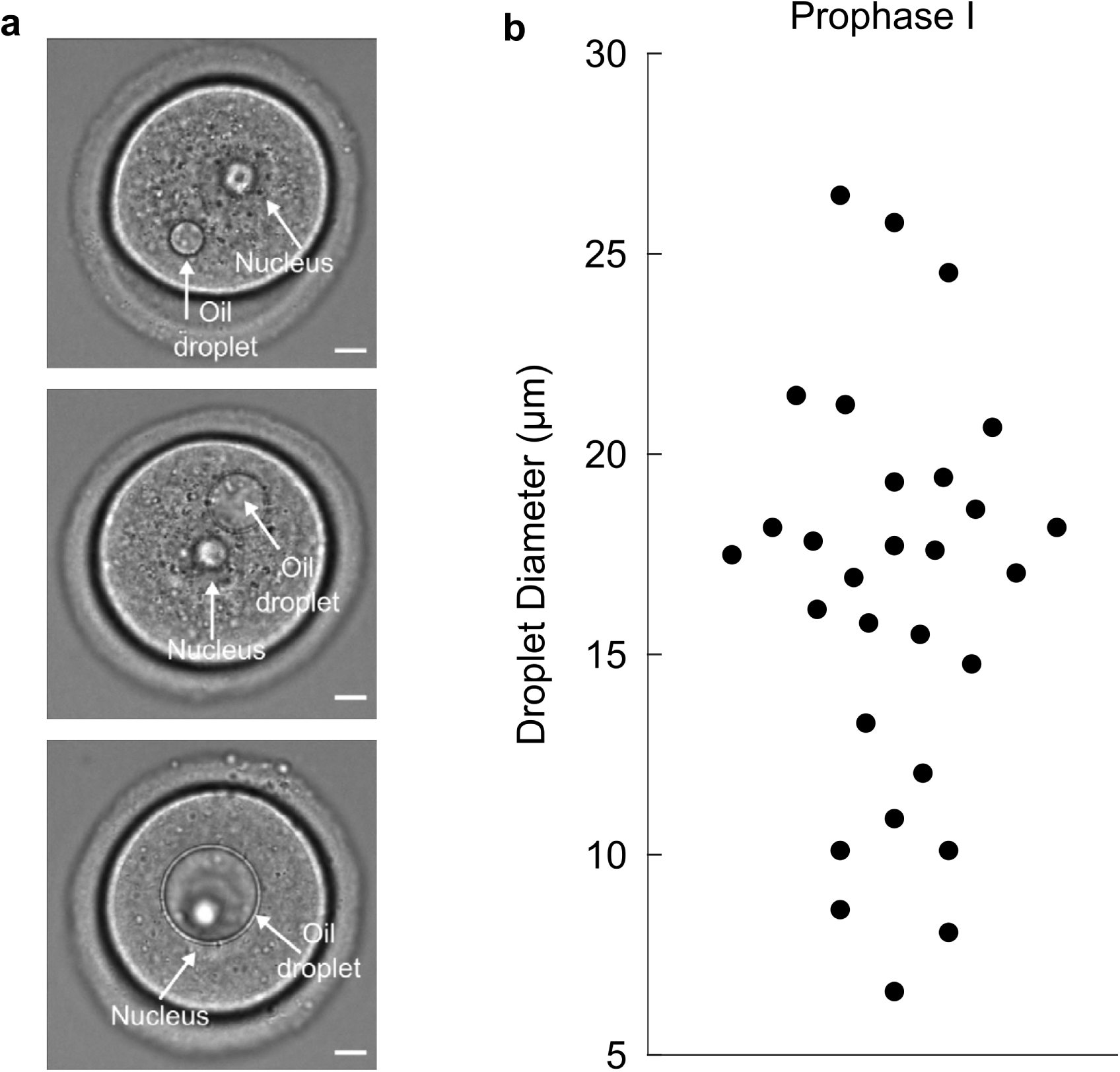
Typical examples of oil droplets of three different sizes injected into Prophase I mouse oocytes. **(a)** Examples of oil droplets in Prophase I mouse oocytes (transmitted light, top to bottom: smallest to largest oil droplet). Scale bar is 10 µm. **(b)** Distribution of diameter lengths for all droplets that are not central at the beginning of the experiment (n = 29 oocytes).

**Supplementary Figure S4.**
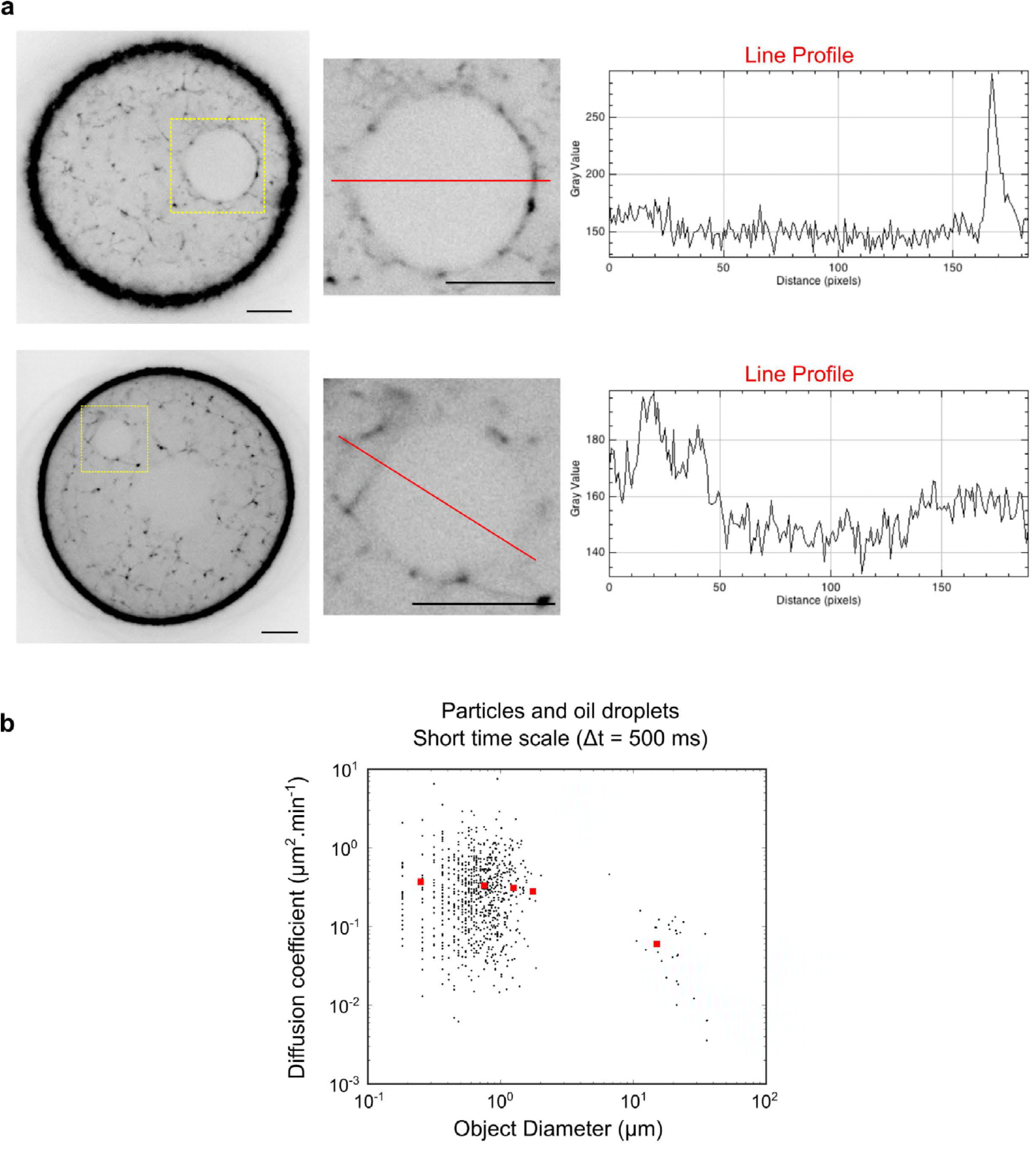
**(a)** Left panels: two examples (up and down) of inhomogeneities of actin recruitment around the droplet after microinjection of GFP-UtrCH. Actin accumulation can be detected on the side facing the cortex compared to the side of the droplet facing the center of the oocyte. F-actin is in black. The yellow dotted squares are presented magnified in the two middle panels. Scale bars are 10 µm. Right panels: Examples of GFP-UtrCH fluorescence intensity levels obtained from the two red line scans highlighted in the middle panels. **(b)** Black points: diffusion coefficients computed from the MSD of the observed objects (particles or oil droplets) at short time scale. Diffusion coefficient was computed from a linear fit on the 20 first points of the MSD curve (10 s). Red squares: diffusion coefficients computed from the distribution of ∆r.cos(θ) at 2.5 s. Diffusion coefficient is defined as: 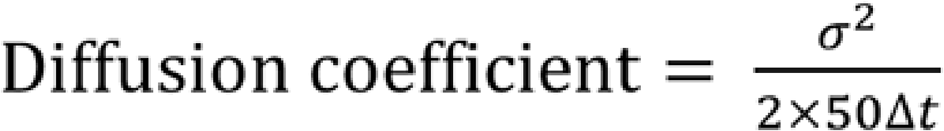 n = 5 oocytes for aggregates of particles, n = 29 oocytes for oil droplets.

**Supplementary Figure S5.**
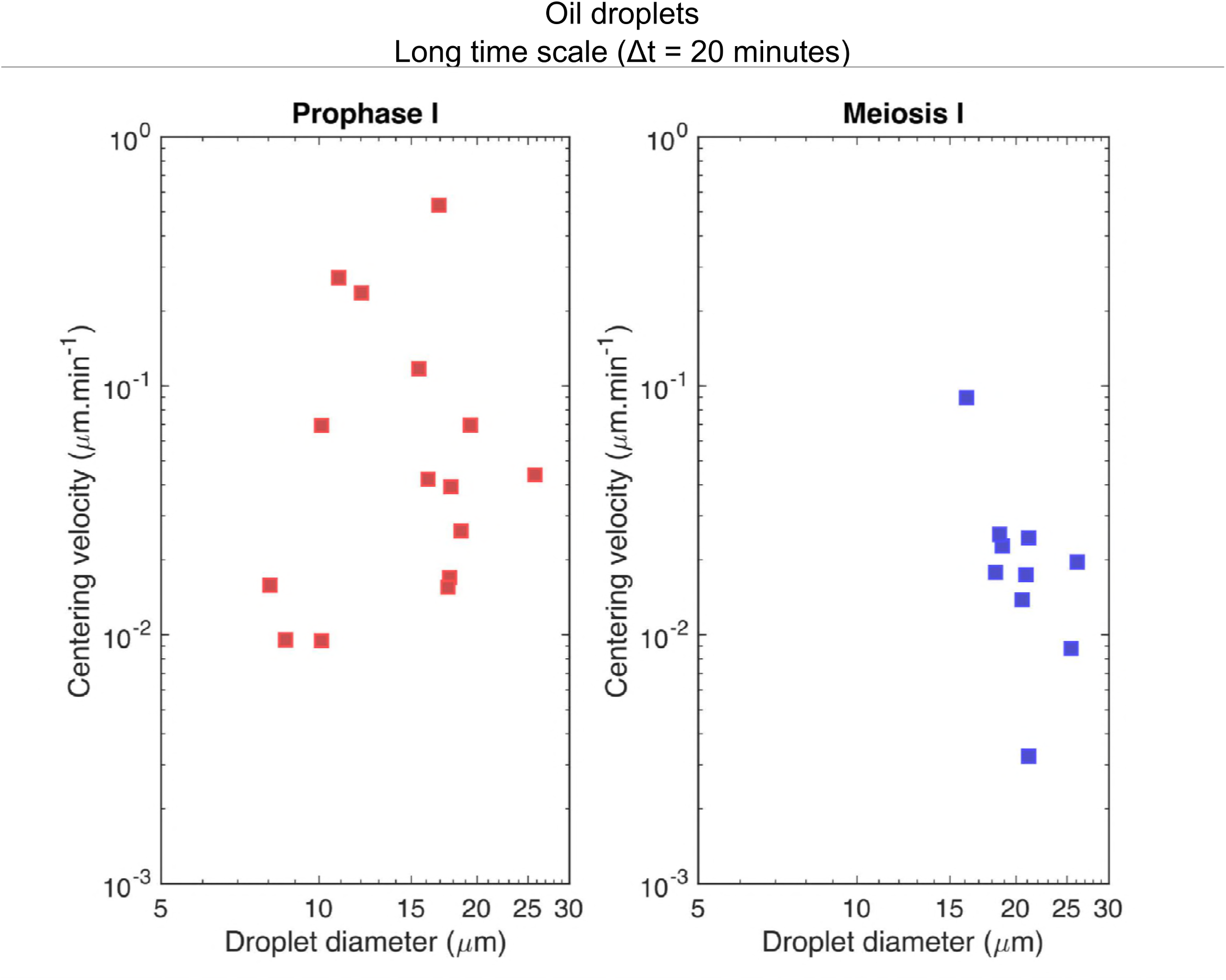
Mean centering velocity as a function of the droplet diameter in Prophase I and Meiosis I. The mean centering velocity is measured as explained in Fig. 4b. n = 22 oocytes in Prophase I and n = 11 oocytes for Meiosis I.

**Supplementary Figure S6.**
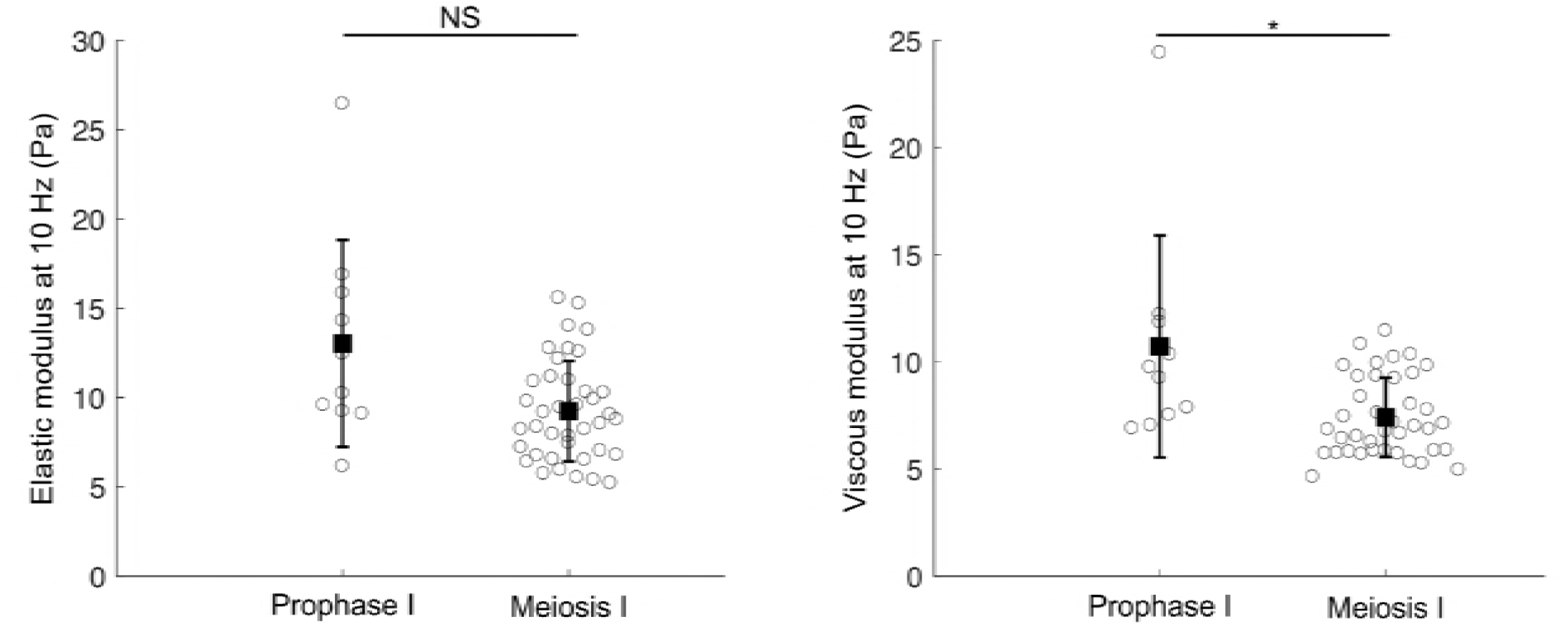
Optical tweezers measurements. Quantification of elastic and viscous moduli at 10 Hz for oocytes maintained in Prophase I and oocytes undergoing Meiosis I. For oocytes undergoing Meiosis I, the measurements were taken at NEBD + 6h. p-values were calculated with a Kolmolgorov Smirnov statistical test (p-value = 0.07 for G’, the elastic modulus on the left panel and p-value = 0.02 for G’’, the viscous modulus on the right panel). n = 10 vesicles in 7 oocytes in Prophase I and n = 39 vesicles in 14 oocytes in Meiosis I. Mean and S.E.M. are superimposed on the raw data.

**Supplementary Figure S7.**
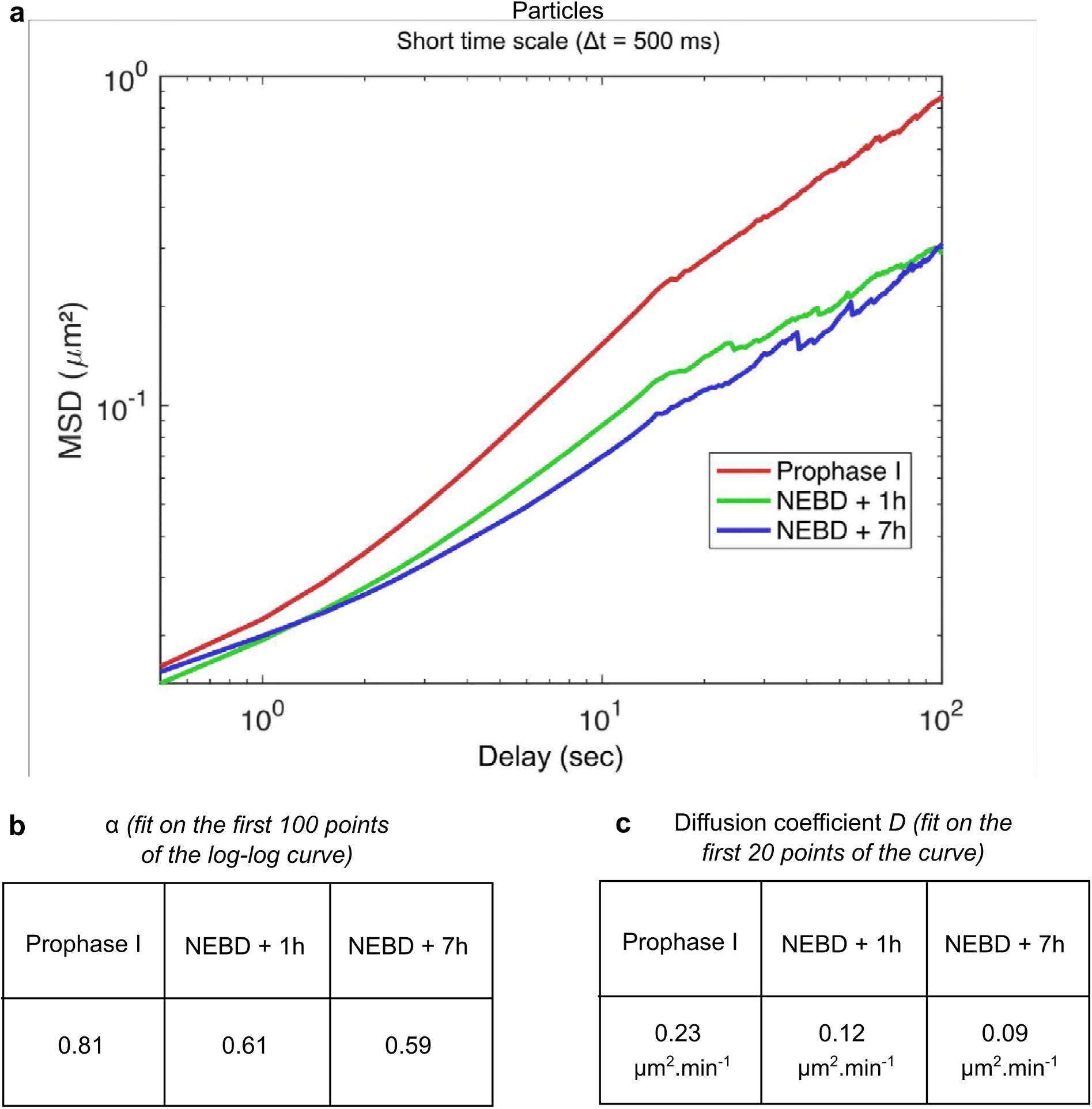
MSD of vesicle colored with Nile Red in Prophase I and Meiosis **I**. **(a)** Log-log representation of the MSD plotted in Fig. 4c right panel. The mean of the MSD for 10 oocytes in Prophase I (red curve) and 12 oocytes at NEBD +1h (green curve) and NEBD + 7h (blue curve) are presented. **(b)** Results from fitting of the MSD in a log-log scale. α represents the slope of the curve (here fit on the first 100 points of the curve). **(c)** Results from fitting of the MSD to extract the diffusion coefficient, D, as done on Fig. S1.

## Supplementary Videos legends

**Supplementary Video 1. Oil droplet in an oocyte maintained in Prophase I at short time scale.** Time lapse movie of an oocyte injected with oil droplet in Prophase I. Frames are taken every 500 ms. Movie duration is 500 s.

**Supplementary Video 2. Oil droplet in an oocyte maintained in Prophase I treated with Cytochalasin D at short time scale.** Time lapse movie of an oocyte injected with oil droplet in Prophase I and treated with Cytochalasin D. Frames are taken every 500 ms. Movie duration is 300 s.

**Supplementary Video 3. Oil droplet in an oocyte maintained in Prophase I at long time scale.** Time lapse movie of an oocyte injected with oil droplet in Prophase I. Frames are taken every 20 min. Movie duration is 16 h.

**Supplementary Video 4. Oil droplet in an oocyte undergoing Meiosis I at long time scale.** Time lapse movie of an oocyte injected with oil droplet and undergoing Meiosis I. Frames are taken every 20 min. Movie duration is 14 h.

**Supplementary Video 5. Oil droplet in an oocyte undergoing Meiosis I and treated with Cytochalasin D at long time scale.** Time lapse movie of an oocyte injected with oil droplet, treated with Cytochalasin D and undergoing Meiosis I. Frames are taken every 20 min. Movie duration is 14 h.

**Supplementary Video 6. Total vesicles observed at different states of meiosis.** Time lapse movies of oocytes stained with Nile red at various states of meiosis (from left to right): Prophase I, Meiosis I early (NEBD+1h) and Meiosis I late (NEBD+6h). Frames are taken every 500 ms. Movie duration is 150 s.

